# The hemodynamic initial-dip consists of both volumetric and oxymetric changes correlated to localized spiking activity

**DOI:** 10.1101/259895

**Authors:** Ali Danish Zaidi, Niels Birbaumer, Eberhard Fetz, Nikos Logothetis, Ranganatha Sitaram

## Abstract

The “initial-dip” is a transient decrease frequently observed in functional neuroimaging signals, immediately after stimulus onset, and is believed to originate from a rise in deoxy-hemoglobin (HbR) caused by local neural activity. It has been shown to be more spatially specific than the hemodynamic response, and is believed to represent focal neuronal activity. However, despite being observed in various neuroimaging modalities (such as fMRI, fNIRS, etc), its origins are disputed and its neuronal correlates unknown. Here, we show that the initial-dip is dominated by a decrease in total-hemoglobin (HbT). We also find a biphasic response in HbR, with an early decrease and later rebound. However, HbT decreases were always large enough to counter spiking-induced increases in HbR. Moreover, the HbT-dip and HbR-rebound were strongly coupled to highly localized spiking activity. Our results suggest that the HbT-dip helps prevent accumulation of spiking-induced HbR-concentration in capillaries by flushing out HbT, probably by active venule dilation.

## Introduction

Functional neuroimaging is a powerful non-invasive tool for studying brain function in health and disease, that uses changes in blood oxygenation as a proxy for estimating local neuronal activity [1]. However, which feature of the hemodynamic signal best reflects local neuronal activity remains an open question. The most commonly used feature is the hemodynamic response amplitude, which is slow and unspecific[1]. Since neuronal processes such as multi-unit spiking are fast, dynamic and spatially localized, a feature in the BOLD signal with similar properties, which also correlates strongly with local neuronal activity, would be an ideal candidate. Early fMRI studies reported such a quick and localized dip in the initial BOLD signal immediately following stimulus onset in various brain areas[2], [3]. This early decrease was termed the ‘initial-dip’, and was believed to originate from a rise in deoxy-hemoglobin (HbR) caused by stimulus-induced changes in localized neuronal activity[2]. Supporting evidence comes from studies reporting spatially localized dips in tissue partial oxygen pressure[4], [5], and increases in the concentration of HbR [6], observed at the time of the dip. The initial-dip is also more spatially localized than the positive BOLD response [7], and has been used to accurately map orientation columns in the visual cortex (better than the positive-response)[8]. Based on these observations, the initial-dip is believed to represent focal neuronal activity [8]. Although the initial-dip has been observed in various functional neuroimaging modalities (such as BOLD-fMRI [2], [7], [9], optical imaging [10], [11], fNIRS[12] and pO_2_-measurements [4]), its origins are disputed [11] and its precise neuronal correlates are all but entirely unknown.

We recently documented a method for the simultaneous acquisition of epidural fNIRS and intra-cortical electrophysiology in primates (Fig. 1A-B), demonstrating that fNIRS has high SNR when acquired epidurally[12], making it ideal for studying local neuro-vascular interactions. FNIRS uses a light-emitter and detector pair (optode pair) to measure changes in concentrations of oxygenated (HbO), deoxygenated (HbR) and total (HbT) hemoglobin, within the vascular compartments in a small volume of tissue[13], [14]. We recorded runs of both spontaneous and stimulus-induced activity in the primary visual cortex of two anesthetized monkeys.

**Figure 1.**
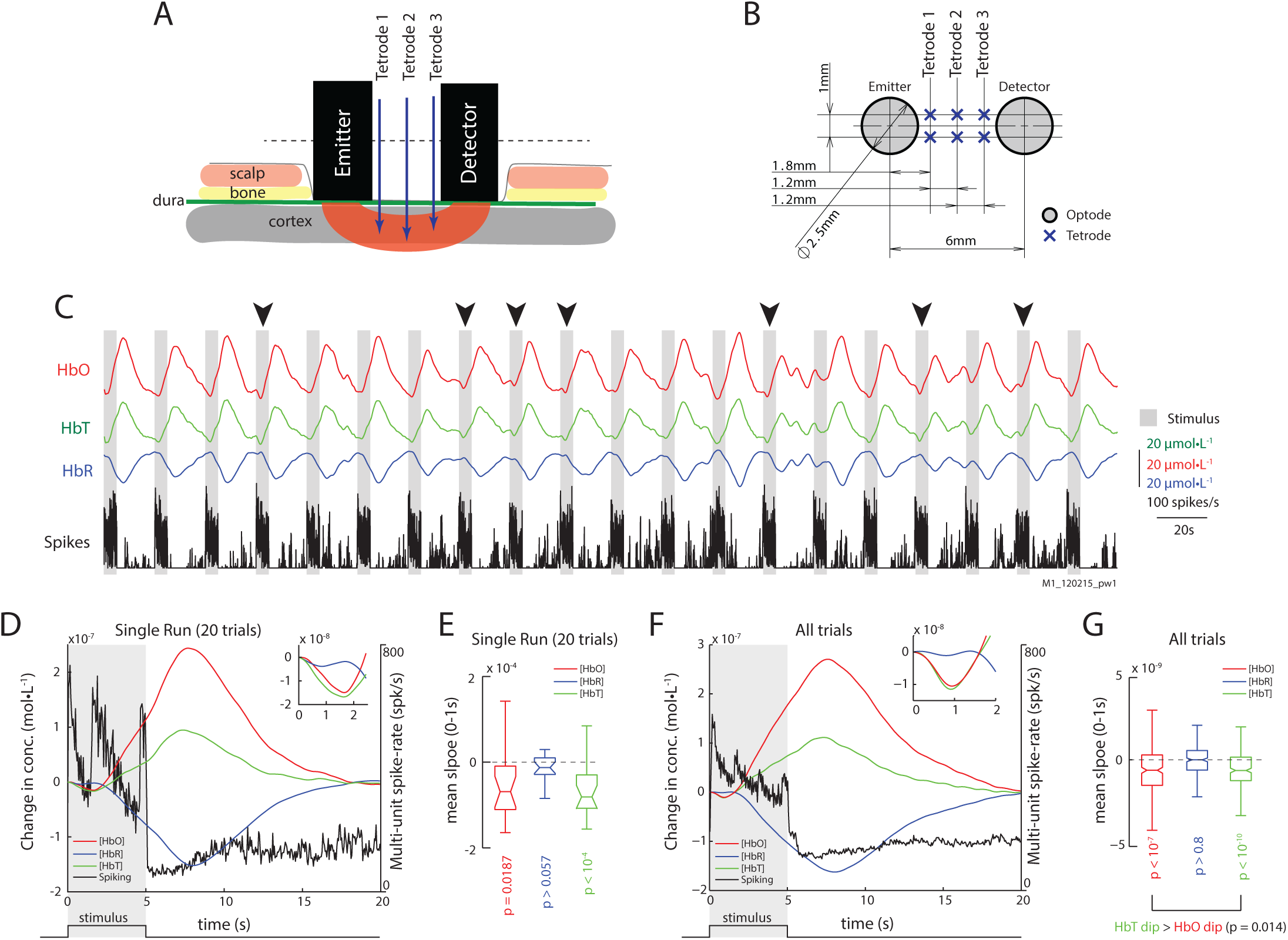
Epidurally measured fNIRS measurements reveal initial dips in hemodynamic signals. A) Illustration of the sensor array with placement of fNIRS optodes and electrodes relative to scalp and brain tissue. B) Transverse section of the sensor array with distances between optodes and electrodes. See methods for details. C) Traces of HbO, HbR, and Spiking from an example run with 20 trials. Grey bars represent epochs of visual stimulation. Arrows mark trials where initial dips are obvious in signal trends. D) The mean traces of HbO, HbR, HbT and multi-unit spiking (units on the right) for trials shown in (C). Inset) Same hemodynamic traces, but from 0-2.5s. The initial dip is observed in the HbO and HbT (inset), but not in the HbR traces. The shaded region represents visual stimulus presentation. E) Distribution of slopes from 0-1s for HbO, HbR and HbT traces for trials in (C). The distributions of HbO and HbT slopes are less than zero, but not for those for HbR (p_HbO_=0.0187; p_HbT_<10^−4^; p_HbR_=0.099; n=20; Š). F) The mean traces of HbO, HbR, HbT and multi-unit spiking activity (units on the right) for all trials. Inset) Same hemodynamic traces, but from 0-2s. G) Distribution of signal slopes from 0-1s for HbO, HbR and HbT traces for all trials. The distributions for HbO and HbT are less than zero, but not for HbR (p_HbO_<10^−7^; p_HbT_<10^−10^; P_HbR_>0.1; n=260; Š) However, HbT dips were stronger than HbO dips (p = 0.028; Ś)

## Results

Fig. 1C shows the traces of HbO, HbR, HbT and multi-unit spike-rates for an example run with visual stimulation, consisting of 20 trials. The grey bars mark the 5s of visual stimulation (whole-field rotating chequerboard with high contrast), which were followed by 15 seconds of rest (white spaces). Obvious dips in the HbO signals can be observed on some trials (black arrows). The average traces of these 20 trials elicited observable dips in both HbO and HbT (Fig. 1D). We used the mean signal slope between 0 and 1s as a metric of the ‘strength’ of the initial-dip for each hemodynamic signal, and found that both the HbO and HbT traces had significant dips (Fig. 1E). Similarly, in the mean traces of all 260 trials from our dataset, a clear dip in the HbO and HbT signals can be observed, without any changes in HbR (Fig. 1F-G, Table 1).

To understand the relationship between neuronal activity and the initial-dip, we divided the 260 trials in our dataset into two groups based on the peak spike-rate during stimulus onset, namely, high-spiking trials (899.96 ± 12.89 spk/s; n=122) and low-spiking trials (497.28 ± 8.35 spk/s; n=125) (Fig. 2A). For high-spiking trials we observed strong dips in HbO, HbR and HbT traces. Although the overall distribution of dips in HbO and HbT were not significantly different (p>0.32; Wilcoxon’s rank-sum test), a trial-by-trial comparison revealed that HbT dips were in fact larger (p<10^−5^, Wilcoxon’s signed-rank test; Fig. 2B-C). Although low-spiking trials seem to elicit faint modulations in the HbO and HbT signals, we did not observe significant changes in their corresponding slopes (p<0.1; Fig. 2C). The low-spiking trials themselves, however, had both significantly high peak spike-rates, and strong stimulus-induced spike-rate modulations (Fig. 2D). Furthermore, in high-spiking trials, we observed a biphasic response in the slope of HbR traces, which was absent in the low-spiking trials (Fig. 2E). The HbR elicited a small but significant dip (between 0-0.75s, epoch I) and a later rebound in the high spiking trials (defined as mean HbR slope between 0.75-1.75s, epoch II, Fig. 2F). This illustrates that there is indeed an increase in HbR signal with higher spiking activity.

**Figure 2.**
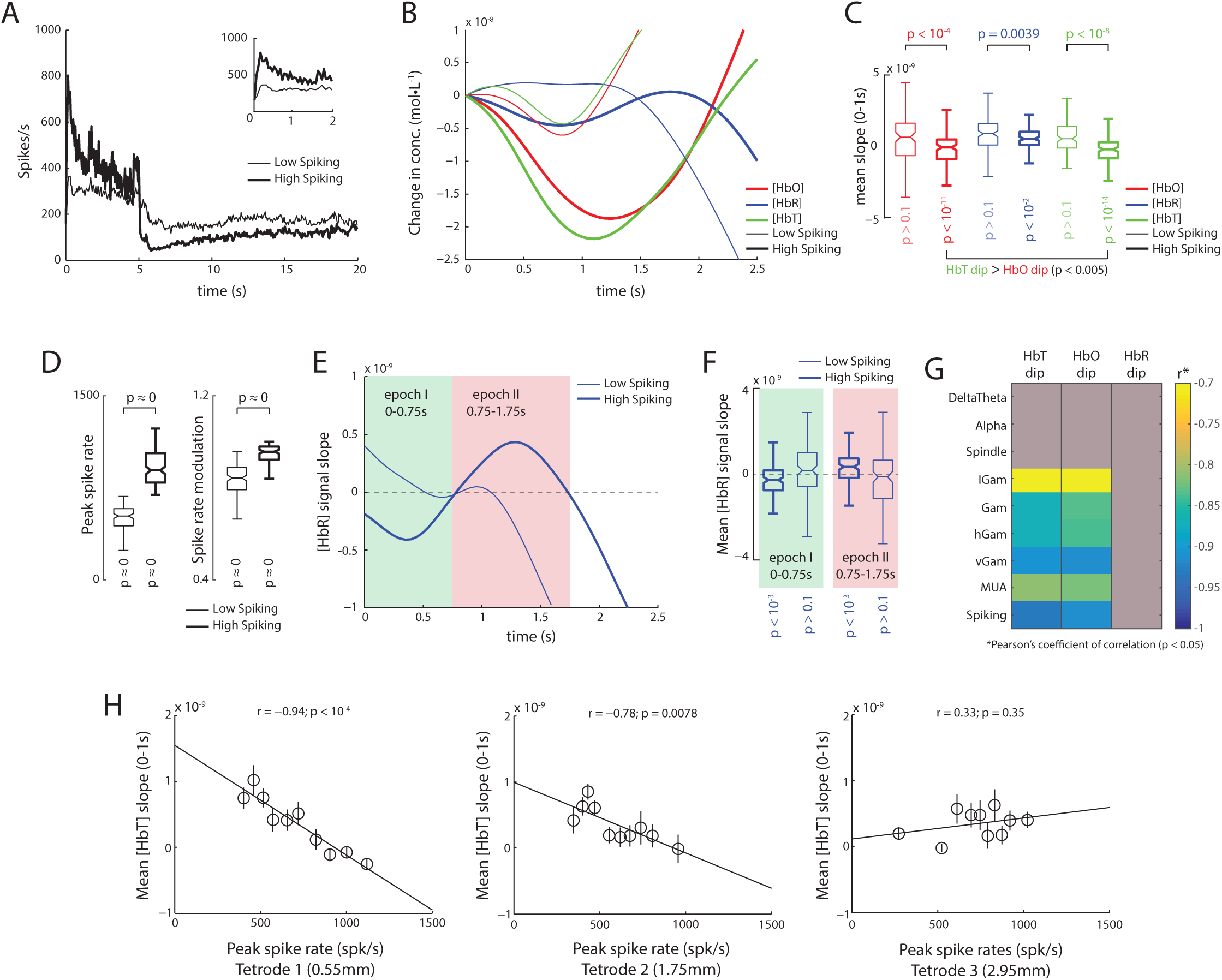
Trials with high spiking activity reveal initial dips comprise of an early HbT decrease, and late HbR increase. A) Mean traces of spike-rates for trials with high and low spiking immediately after stimulus onset (thick and thin traces, respectively). Inset) Same traces, but between 0-2s. B) Mean traces of hemodynamic signals for trials with low (thin) and high spiking as shown in (A). A clear increase in the dips is observed for high spiking trials with the largest dips elicited in HbT traces. C) Average slopes from 0-1s for HbO, HbR and HbT traces for high (thick) and low (thin) spiking trials. HbO, HbR and HbT all elicit significant dips in high-spiking trials (p_HbO_<10^−11^; p_HbT_<10-^14^; p_HbR_<10^−2^; n=128; Š), with larger dips in HbT than HbO (p<0.005; n=125; pairwise Š). Interestingly, trials with low spiking trials do not elicit significant dips in either HbO, HbR or HbT (p<0.1; n=122; Š). D) Distribution of peak spike-rates and visual modulation of spike-rates for trials with high (thick) and low (thin) peak spike-rates. This illustrates that even though the peak rates were lower in the low-spiking trials, the overall spiking activity was significantly high, as was the visual stimulus induced modulation of spike rates (see Methods for calculation of modulation index). E) Analysis of the slope of HbR traces in high spiking trials reveals a biphasic response, which is almost all but absent in low spiking trials. In high spiking trials (thick trace), an initial negative slope is observed roughly between 0-0.75s (epoch I, shaded green), followed a positive slope roughly between 0.75-1.75s (epoch II, shaded red). F) For high spiking trials, the distribution of mean HbR slopes were significantly negative during epoch l (p<10^−3^; n=125; Š), and significantly positive during epoch II (p<10^−3^; n=125; Š). In contrast, low spiking trials showed no significant modulation of HbR slopes during either epoch I or II (p>0.1; n=122; Š). G) Correlation of mean dips in HbT, HbO and HbR stimulus induced peaks in the power of various LFP frequencies bands and Spiking. Correlations with p>0.05 are greyed. Only high frequency bands showed a significant correlation with initial dips, with spiking activity eliciting the strongest relationship, that were marginally higher for HbT than HbO. H) Strength of the relationship between the HbT dip and spiking activity decreases with distance from the NIRS emitter. Strongest correlations are observed on tetrode closest to emitter (0.55mm away from emitter edge, 1.8mm from emitter center), whereas no correlations are observed on tetrode 2.95mm away (4.2mm from center).

We next assessed the relationship of the initial-dip with various bands of the local field potential (LFP). We filtered the extracellular field potential into eight frequency bands, namely the DeltaTheta (1-8 Hz), Alpha (9-15 Hz), Spindle (15-20 Hz), low-Gamma (lGam, 20-40 Hz), Gamma (Gam, 40-60 Hz), high-Gamma (hGam, 60-100 Hz), very high-Gamma (vGam, 125-300 Hz) and multi-unit activity (MUA, 1-3 kHz) bands, and obtained their respective band envelopes (see Methods for details). From the various LFP bands we analyzed, only peaks in high-frequency bands had a significant correlation with the HbO and HbT dips (Fig. 2G). However, the strongest dependence was still observed with peaks in spiking activity for both HbO and HbT dips, with slightly higher correlations observed with HbT than HbO (Fig. 2G, Fig. S1). We next determined how this relationship with spiking varied as a function of distance over cortical surface. We obtained the correlations between the HbT dip and the peak spike-rates on the three tetrodes placed between the emitter and detector. We found that the correlation was strongest with the tetrode closest to the emitter, and that it decreased with increasing distance from the emitter (r = −0.94; p <10^−4^, Fig. 2H). The results were identical when we used the peak-amplitude of the initial-dip instead of the mean signal slope for both HbO and HbT dips (Fig. S2). This finding not only corroborates the idea that the initial-dip is a spatially localized hemodynamic response, but also provides neurovascular evidence that fNIRS has a spatial sampling bias in favor of the emitter, questioning the popular “banana model” that assumes uniform sampling through the volume of tissue between the emitter and detector[13].

It might be argued that stimulus induced activity introduces artificial correlations between the neuronal and hemodynamic responses, by inducing highly synchronous spatio-temporal patterns [15]. To ensure our results didn’t arise from such correlations, we analyzed recordings of spontaneous ongoing activity in the absence of visual stimulation, where the monkeys’ eyes were closed and covered with thick gauze. Fig. 3A shows traces of HbO, HbR, HbT and spike-rates for an example run of spontaneous activity, consisting of 15 minutes. Dips in the HbO and HbT signals can be seen to coincide with strong bursts in spiking activity (Fig. 3A, arrows and bars). To analyze the relationship between spiking and hemodynamic signals, we used system identification to estimate the impulse response from spiking to HbO, HbR and HbT traces. This method uses the Wiener-Hopf relationship [16] to estimate the impulse-response to a unit-pulse (1 SDU amplitude, 1s duration at t=0) from the input (spiking activity) on the output (hemodynamic signal), and is independent of the shape and auto-correlation structure of the input. Fig. 3B shows the mean impulse-responses from spiking on HbO, HbR and HbT traces. There is an evident dip in the HbO and HbT traces. However, the slopes of these impulse response functions provide a clearer picture of the signal dynamics (Fig. 3C), where a decrease in HbO and HbT can be observed at t=1s (Fig. 3D; n=48; **Š**), and a late rebound of HbR can be observed at t=2s. We next used the total spike count in each run to separate the runs into high-spiking and low-spiking runs (Fig. 3E). Fig. 3F shows the impulse-responses obtained for the high-spiking (thick) and low-spiking (thin) runs. The slopes of the impulse-responses show that the high-spiking trials had large, significant dips for all three hemodynamic signals (HbO, HbR and HbT; Fig. 3G-H). Furthermore, we found no significant difference between the overall distributions of HbO and HbT dips (p>0.45, n=85, Wilcoxon’s rank-sum test). However, on trial-by-trial comparison, HbT dips were significantly stronger than HbO-dips (Fig. 3H; p<10^−4^, n=85, Wilcoxon’s one-tailed signed-rank test). Interestingly, although the low spiking runs seemed to elicit dips as well, they did not reach significance. Furthermore, only the high-spiking runs elicited significant modulations in HbR with both significant dips at t=1s and rebounds at t=2s (Fig. 3I). These results are identical to those obtained from the analysis of stimulus induced activity, and are thus independent of the visual stimulation paradigm.

**Figure 3.**
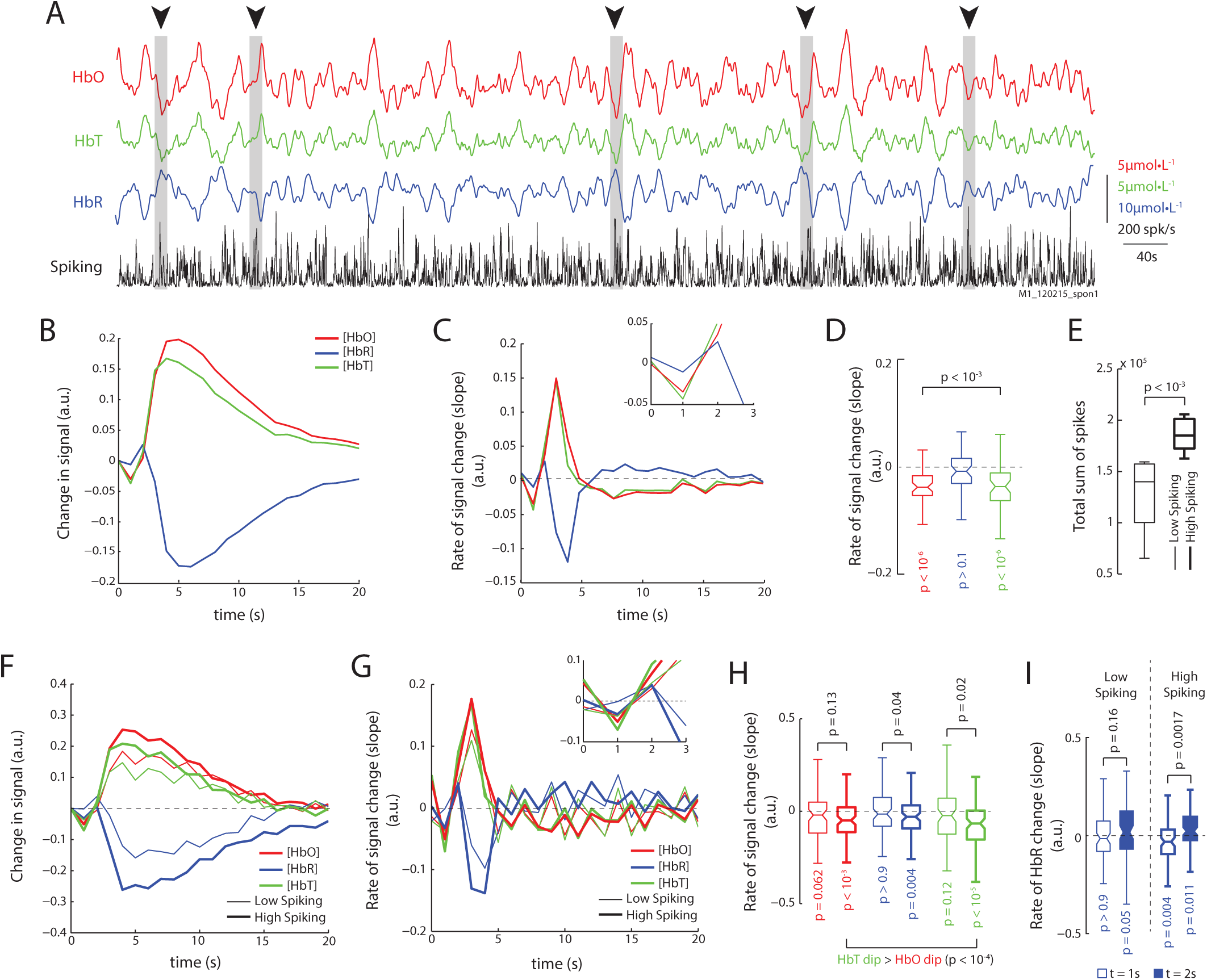
Analysis of spontaneous activity in the absence of visual Stimulation reveals identical relationships. A) Traces of HbO, HbR, HbT and spike-rates from an example run of 900s. Periods of high spiking activity that elicit an observable dip in HbO and HbT are marked with arrows and grey bars. B) We used system identification to estimate the impulse response functions from spiking to HbO, HbR and HbT signals in recordings of spontaneous activity. The mean impulse response reveals a dip in HbO and HbT (mean of 48 impulse response functions obtained from 16 runs lasting 900s each; see Methods for details). C) Rate of change of the impulse response functions for HbO, HbR and HbT reveals dips in both HbO and HbT, and a late rise in the HbR. Inset) Same traces but between 0 to 3s. D) Distribution of slopes for HbO, HbR and HbT at t=1s. Only HbO and HbT have significant dips, but not HbR (Š; n=48). E) The runs were divided based on the total sum of spikes in each run, and seperated into low spiking and high spiking runs. High spiking runs had signficanlty higher spike sums (Š; n=8). F) The mean impulse responses for high and low spiking runs reveal stronger modulation of hemodynamic signals on high spiking trials. Color-code same in following figures G) Mean traces of slopes of impulse responses shown in F. High spiking trials elicit an obvious dip at t=1s. Inset) Same traces but from 0 to 3s. H) Distribution of dips for low and hihg spiking trials (legend same as F). Only high spiking trials have significant dips in all three signlas. Also, HbT dips were larger than HbO dips (Š; n=80). I) When comparing the HbR dip and rebound at t=1s and t=2s resp., only high spiking trials reveal a strong dip and rebound in the HbR signal.

In the analysis of both spontaneous and stimulus-induced activity, we find that the initial-dip is dominated by a decrease in HbT, in trials with strong bursts in spike-rates. However, it might be argued that this decrease in HbT is not an actual change in blood volume, but a consequence arising from signal trends, such as the slope of the hemodynamic signal before stimulus onset, or the choice of analytical parameters, such as the differential path-length factors (DPFs) used for the conversion of optical density to concentration changes (the only parameter-dependent transformation in the analysis). Surprisingly, the strength of the initial-dip failed to correlate with the mean slope of the hemodynamic signal 0-2s prior to stimulus onset (r=-0.002, p>0.9, Pearson’s correlation coefficient of correlation), suggesting that the trend of the hemodynamic signal before the dip fails to affect the initial-dip. We also used various combinations of physiologically relevant DPFs, as reported earlier[6], in the estimation of concentration changes in HbO, HbR and HbT, and obtained identical results (see Fig. S3). To be sure, we also analyzed the raw optical density changes for both wavelengths (760nm and 850nm). We found a significant decrease in the optical densities for both chromophores, which was enhanced in high-spiking trials (see Fig. S4). The decrease in chromophore concentration at both wavelengths can only be attributed to decreases in total hemoglobin concentration (HbT). Furthermore, within trials with low spiking activity, even though we do not see significant changes in the slope of the HbO and HbR signals, we do find small but significant changes in their concentration (see Fig. S5). We detected significant increases in the HbR concentration (mean HbR concentration change between 0 to 0.8s, p<0.05, Wilcoxon’s signed-rank test) as well as significant decreases in the HbO concentration (see Fig. S5C, p < 0.05; n=125; Wilcoxon’s sign-rank test) but failed to detect significant changes in HbT (p>0.1, n=125; Wilcoxon’s signed rank test), an observation that is in agreement with previous reports on the initial-dip [6], [17]. Finally, the trials within the lowest quartile of spike-rates elicited neither initial-dips (in HbO, HbR or HbT traces, mean slope between 0-1s), nor changes in concentration (mean concentration change between 0-1s), even though these trials still had significantly high bursts in spike rates (peak rate 300±40 spk/s; p<10^−11^; n=59, Wilcoxon’s signed-rank test). These observations demonstrate two different manifestations of the initial-dip. During low-spiking, there is an oxymetric change consisting of an increase in HbR and decrease in HbO concentration. During very high spiking, there is a volumetric change, consisting of a decrease in HbT (and consequently HbO and HbR).

Interestingly, all significant dips detected in optical density traces also translated to changes in hemoglobin concentration, irrespective of the choice of parameters used for converting optical density to concentration change (**see** Fig. S3). Combined with previous studies [6], these results are in conflict with an earlier report where changes in optical density failed to translate to changes in HbO, HbR or HbT signals[11].

## Discussion

Although the exact vascular compartments that fNIRS samples from have not yet been firmly established, it is generally believed to reflect oxymetric changes within vessels smaller than 1mm in diameter[14], such as pre-capillary arterioles, capillaries and post-capillary venules (Capillary and Peri-capillary Vessels, henceforth CPVs), which is where most of the oxygen-exchange occurs[18], and hence where the largest changes in blood oxygenation occur. This is exemplified by the increase in HbO and decrease in HbR observed during influx of oxy-saturated blood into CPVs during the positive hemodynamic response (Fig. 1F). Thus, even though neuronal activity leads to an increase in HbR concentration, the HbR signal during the positive response decreases, once oxy-saturated blood passes through the CPVs post arteriole dilation. Even the transient increases observed in the HbR signal during the dip only last until the HbT response commences. Hence, a decrease in both HbO and HbR during the initial-dip most likely represents decreases in the blood-volume within CPVs. In our data, the HbO/HbT dip ratio (the ratio of HbO to HbT decrease at maximal dip) is 50.4±17% (mean±sem) for stimulus-induced, and 58.9%±32% for spontaneous activity, which is within the range of oxygen saturation reported within CPVs [18]. A possible means to attain a decrease in CPV blood volume would be through the dilation of post-capillary venules. Post-capillary venules have recently been shown to have band-like smooth muscles encircling their circumference, similar to those associated with precapillary arterioles [19]. Furthermore, small venules have been shown to increase their diameter simultaneously with strong arteriolar dilations in recordings of spontaneous activity [20]. However, since the primary arterioles that are dilated are further away from the capillaries, the influx of blood takes longer to reach the capillaries. Therefore, upon post-capillary venule dilation, the decrease in capillary blood pressure would briefly compress the capillary, flushing the blood out, before the influx of oxygenated blood caused by arteriole dilation. Recently, erythrocytes have been reported to deform with reduced oxygen tension, facilitating an increase in their flow-rate through the capillary lumen[21], aiding this process. Indeed, a transient increase in capillary RBC velocity can be observed immediately after stimulus onset, which briefly subsides, before finally increasing again (see figures 4B and 4D in 20).

Such a flushing of CPV blood could serve to prevent HbR accumulation in the capillaries, enforcing a virtual “upper-limit” of HbR concentration in the vascular tissue, as well as facilitating the influx of oxygen saturated blood from the arterioles. Indeed, in our data, we found that spiking correlated strongly with the HbR-rebound (Fig. 4A). However, the HbT dips were consistently larger than the HbR-rebounds, maintaining an upper-limit of HbR concentration, and hence no relationship was observed between spiking and HbR concentration change between 0.75-1.75s (Fig. 4B). Moreover, when corrected for the HbT dip (by subtracting HbT traces from HbR traces), the “dip-corrected” HbR traces reveal increases in concentration that are correlated to spiking activity (Fig. 4C). In contrast, no such relationship is observed with “dip-corrected” HbO traces (Fig. 4D), illustrating that the initial-dip counters rising HbR concentration in the vascular tissue. This deoxygenated blood flushed from individual CPVs would flow into the surface venules, transiently increasing their HbR concentration. Indeed, cortical-depth resolved BOLD-fMRI, believed to reflect changes in tissue HbR [1], [22], reveals that the amplitude of the initial-dip is largest near the cortical surface, in both human [9] and animal [10] studies. Further experiments quantifying changes in HbO and HbR in the various vascular compartments could shed further light on the exact vascular mechanisms of the initial-dip. Nevertheless, our results conclusively demonstrate that the initial-dips in both HbO and HbT traces are strongly correlated with highly localized spiking activity. Furthermore, since we find no relationships between the initial-dip and low-frequency LFP activity, it establishes that the initial-dip is a highly specific marker of localized bursts in spiking activity.

**Figure 4.**
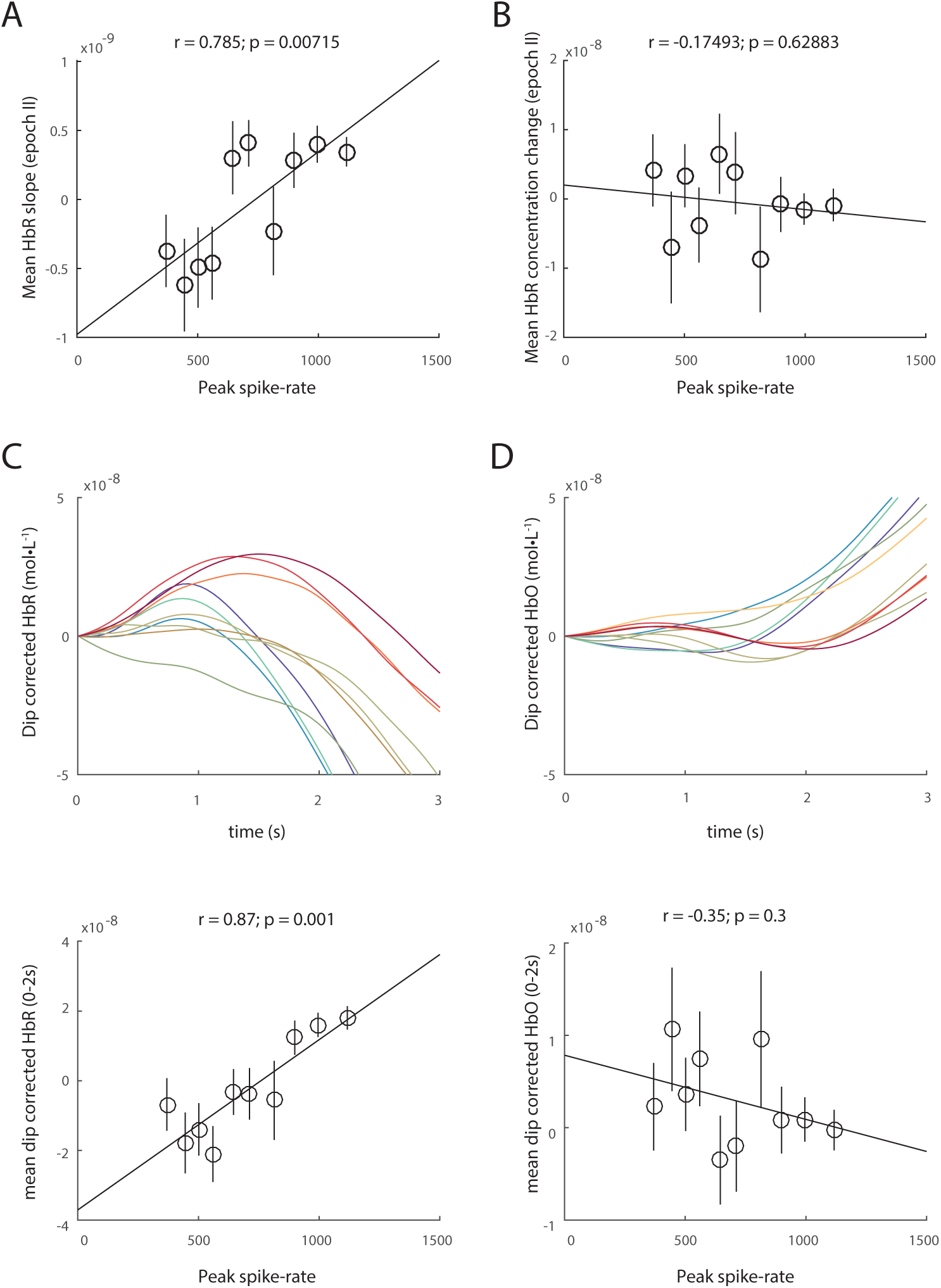
HbR-rebound does not lead to increase in HbR concentration. Although there is a significant correlation between spiking and the mean HbR slope in epoch II (A), the relative HbR concentration change remains unchanged with spiking (B). C) Dip-corrected HbR traces, obtained by imply subtracting the HbT traces from HbR reveals obvious increases in HbR concentration that correlate with spiking. However, no such relationship is observed with dip-corrected HbO traces.

Although we recorded signals from anesthetized monkeys, it has been shown that this anesthesia regime does not significantly alter local neuro-vascular coupling [23]. We also find that fNIRS represents focal neuro-vascular changes close to the emitter, challenging the generally accepted “banana” model that assumes the cortical volume sampled by fNIRS to be uniformly distributed between the emitter and detector [14]. Concurrently, a study comparing simultaneously recorded fNIRS and fMRI signals in humans finds that the voxels correlating best with HbO/HbR changes are consistently closer to the emitter (see Fig. 2 and Table 2 in [24]), though this is not explicitly stated in their results. Overall these results shed further light on the neuro-vascular changes underlying the initial-dip, and enable a better interpretation of functional neuroimaging signals.

In conclusion, we show that the initial dip, though present in both HbO and HbT signals, is dominated by HbT changes that are correlated to highly-localized spiking, demonstrating that these changes are specific to excitatory neuronal activity. This study is, to the best of our knowledge, the first report of an exclusive marker of spiking activity in hemodynamic signals.

## Abbreviations

BOLD: blood-oxygen level dependent signal
CPVs: Capillary and Peri-capillary Vessels (precapillary-arterioles, capillaries and post-capillary venules)
fMRI: functional Magnetic Resonance Imaging
fNIRS: functional Near Infra-Red Spectroscopy
HbO: concentration of oxy-hemoglobin
HbR: concentration of deoxy-hemoglobin
HbT: concentration of total hemoglobin
LFP: local field potential
R: Wilcoxon’s one-tailed rank-sum test
Š: Wilcoxon’s two-tailed signed rank test
T_1_: Tetrode 1 (0.55mm from emitter)
T_2_: Tetrode 2 (1.75mm from emitter)
T_3_: Tetrode 2 (2.95mm from emitter)

## Acknowledgements

We would like to thank Matthias Munk, Cristina Risueno and Rebekka Bernard for help during the collection of data, and Mirsat Memej and Eduard Krampe for their help during the design and construction of the recording system, and Vishal Kapoor and Mastaka Watanabe for feedback on an earlier version of the manuscript. We acknowledge funding from the DFG, CIN and Max-Planck Society.

## Methods

### Surgery and craniotomy

Two healthy adult monkeys, M1 (female; 8 kg) and M2 (male; 10 kg), were used for the experiments. All vital parameters were monitored during anesthesia. After sedation of the animals using ketamine (15 mg/kg), anesthesia was initiated with fentanyl (31 μg/kg), thiopental (5 mg/kg), and succinylcholine chloride (3 mg/kg), and then the animals were intubated and ventilated. A Servo Ventilator 900C (Siemens, Germany) was used for ventilation, with respiration parameters adjusted to each animal’s age and weight. Anesthesia was maintained using remifentanil (0.2-1 μg/kg/min) and mivacurium chloride (4-7 mg/kg/h). An iso-osmotic solution (Jonosteril, Fresenius Kabi, Germany) was infused at a rate of 10 ml/kg/h. During the entire experiment, each animal’s body temperature was maintained between 38.5 °C and 39.5 °C, and SpO2 was maintained above 95%. Under anesthesia, a craniotomy was made on the left hemisphere of the skull to access the primary visual cortex. During each experiment, the bone was removed to create a rectangular slit measuring 3 mm anterio-posteriorly and 20 mm medio-laterally, exposing the dura. Connective tissue, if present above the dura, was carefully removed. For each monkey, at least two weeks were allowed to pass between successive experiments. All protocols were approved by the local authorities (Regierungspräsidium, Tübingen) and are in agreement with European guidelines for the ethical treatment of laboratory animals.

### Near-infrared Spectroscopy

We used a NIRScout machine purchased from NIRx Medizintechnik GmbH, Berlin. The system performs dual wavelength LED light-based spectroscopic measurements. The wavelengths used were 760nm and 850nm, with a maximum of 5μW effective power at the emitter end. Sampling was performed at 20Hz. We used modified emitters and detectors, and optical fiber bundles for sending the light from the LED source into the tissue, and also for detecting refracted light from the tissue. The fiber bundles were ordered from NIRx Medizintechnik GmbH, Berlin, Germany. Both the emitter and detector fiber bundles had iron ferrule tips with an aperture of 2.5mm on the ends that touched the dura. We used three optodes in a linear arrangement separated by 6mm each. Three tetrodes and single-wire electrodes each were added between each pair of adjacent optodes. We used the central optode as a constant detector, and alternated the peripheral optodes during sessions, such that on a given experimental day, 50% of data came from one emitter-detector pair and 50% from the other. The recording instrument was connected via USB to a laptop computer running an interactive software called NIRStar provided along with the instrument. The software was used for starting and stopping recordings, and also for setting up the various parameters, such as, the number of sources and detectors, and the sampling rate. The instrument received TTL pulses from the stimulus system and the electrophysiological recording system, for synchronization purposes. The system sent 1ms TTL pulses every 50ms to the recording system that corresponded to light pulses.

### Electrophysiology

Custom built tetrodes and electrodes were used. All tetrodes and single electrodes had impedance values less than 1 M Ω. The impedance of each channel was noted before loading the tetrodes on to the drive, and once again while unloading the tetrode after the experiment, to ensure that the contacts were intact throughout the duration of the experiment. To drive the electrodes into the brain a 64-channel Eckhorn matrix was used (Thomas Recording GmbH, Giessen, Germany). The electrodes were loaded in guide tubes a day before the experiment. On the day of the experiment, the tetrodes were driven using a software interface provided by Thomas Recording GmbH, Giessen, Germany. The output was connected to a speaker and an oscilloscope, with a switch to help cycle between different channels. We advanced electrodes into the cortex one by one until we heard a reliable population response to a rotating checkerboard flickering at 0.5Hz.

### Spontaneous activity

For each run, spontaneous activity was recorded for 15 minutes, in the absence of visual stimulation. The eyes of the monkey were closed and thick cotton gauze was used to cover the eyelids.

### Visual stimulation

A fundus camera was used to locate the fovea for each eye. For presenting visual stimulation, a fiber optic system (Avotec, Silent Vision, USA) was positioned in front of each eye, so as to be centered on the fovea. To adjust the plane of focus, contact lenses (hard PMMA lenses, Wöhlk, Kiel, Germany) were inserted to the monkey’s eyes. We used whole-field, rotating chequerboard to drive the neural activity. The direction of rotation was reversed every second. Each trial consisted of 5 seconds of visual stimulation followed by 15 seconds of a dark screen. A single run consisted of 20 trials. Data presented are from 13 runs spread over 8 experimental days.

### Signal processing and data analysis

All analyses were performed in MATLAB using custom written code. Only runs that cleared visual screening for artifacts were used. Data points that were larger than 5 SDU were excluded from the analysis, so as to avoid tail-effects for correlation analysis. Normality for each distribution was confirmed before analysis was performed.

### FNIRS signal processing

The raw wavelength absorption data from the NIRS system was converted to concentration changes of [HbO] and [HbR] using a modified Beer-Lambert equation (DPF = 6,6). For correlating hemodynamic signals with neural activity, the signals were filtered between 0.01 and 1 Hz to remove any low frequency drifts. For a trial-by-trial analysis, the hemodynamic response for each trial was zero-corrected by subtracting, from each hemodynamic response, the value at the start of the trial.

### Electrophysiological signal processing

The extracellular field potential signal was recorded at 20.8333 kHz and digitized using a 16-bit AD converted. From the raw signal, eight frequency bands (namely, DeltaTheta (1-8 Hz), Alpha (9-15 Hz), Spindle (15-20 Hz), low Gamma (20-40 Hz), Gamma (40-60 Hz), high Gamma (60-120 Hz), very high Gamma (120-250 Hz) and MUA (1-3 kHz)) were band-pass filtered using a 10^th^ order Butterworth filter. The envelope for each band was then obtained by taking the absolute value of the Hilbert transform of the filtered signal. The band-envelope was then converted to standard deviation units by subtracting the mean and dividing by the standard deviation of the signal. This signal was then resampled at 20 Hz, to allow comparisons with hemodynamic signals.

Spike rates were obtained by detecting peaks in the MUA signal larger than a threshold (2 SDU), and by counting the threshold-crossing events in 50ms bins. Varying the detection threshold between 2, 3 or 4 SDU did not affect the results.

### System identification based impulse response estimation in spontaneous activity

Impulse response functions from spiking to HbO, HbR and HbT were obtained using the system identification toolbox in Matlab. Each hemodynamic signal was first filtered between 0.01 to 1Hz and then normalized by subtracting the mean and dividing by the standard deviation. Spike-counts were obtained by counting the number of threshold crosses (>3 SDU) in the 1-3kHz band in 50ms bins. The bin-count was then divided by the length of the time window to obtain spike rates in spikes/s. Each spike-rate trace was then smoothened by convolving with a Gaussian function of unit amplitude and 100ms standard deviation. The 900s (15 min) recordings were divided into four epochs of 225s each. All signals were then decimated to 1Hz to enable processing with the toolbox.

For estimating the impulse response function for each epoch, the spike-rate for that epoch was used as the input and the hemodynamic signal as output.

### Calculation of modulation indices

The ‘On’ epoch for each trial was defined as the time from 0 to 5.05 seconds. The extra 0.05s were added to accommodate for the off response. The ‘Off’ epoch was defined as the time between 5.05 to 10.05 seconds. The modulation index (MI) was then calculated using the formula: neural modulation = (SR_on_–SR_off_) / (SR_on_+SR_off_); where SR_on_ is the mean spike-rate during Stim On, and SR_off_ is the mean spike-rate during Stim Off for each trial. Runs without significant visual modulation of spike-rates were excluded from the analysis.

### Statistics

All distributions were confirmed to be normally distributed using the Kolmogorov-Smirnov test in Matlab, before using means as a measure of central tendency. All correlation coefficients represent Pearson’s correlation coefficient and corresponding significance values. To calculate the correlations, the trials were sorted and divided into 10 bins (with 26 trials per bin). The mean values of each bin were then correlated. This was done to avoid an otherwise large trial-by-trial variation. The results were independent of the number of bins used for correlation analysis (see Fig. S1)

---

**Figure S1.Related to Figure 1.**
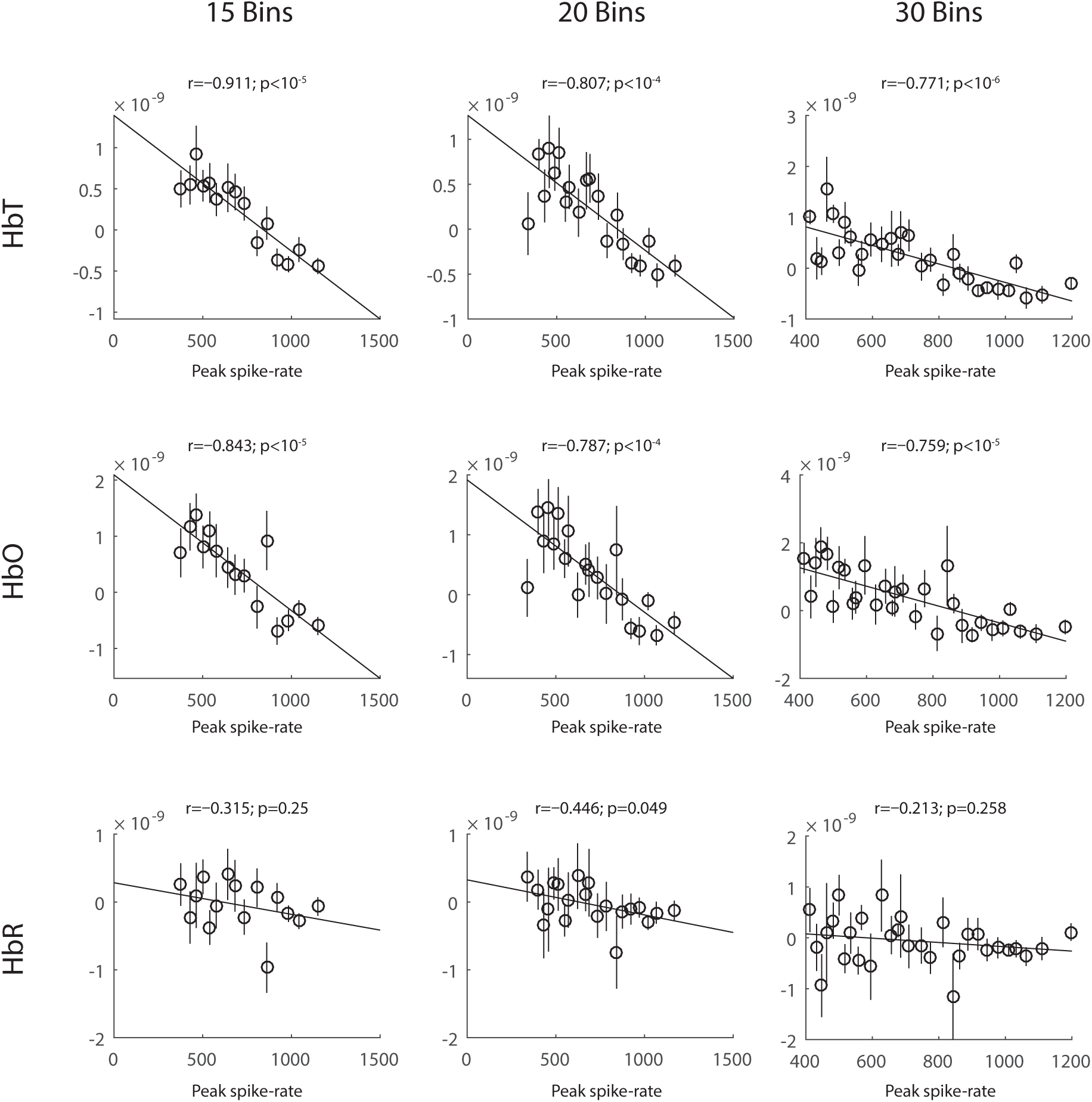
Correlations of spiking activity with HbT, HbO and HbR dips are independent of binsize. Although both HbT and HbO dips showed strong correlations with peaks in spike-rates, the correlations with HbT were stronger than HbO. Although the strength of the correlations got weaker with increasing the number of bins, the significance was not very different. No correlations were observed with the dip strength in HbR. These correlations were independent of the number of groups the data were divided into.

**Figure S2. Related to Figure 1.**
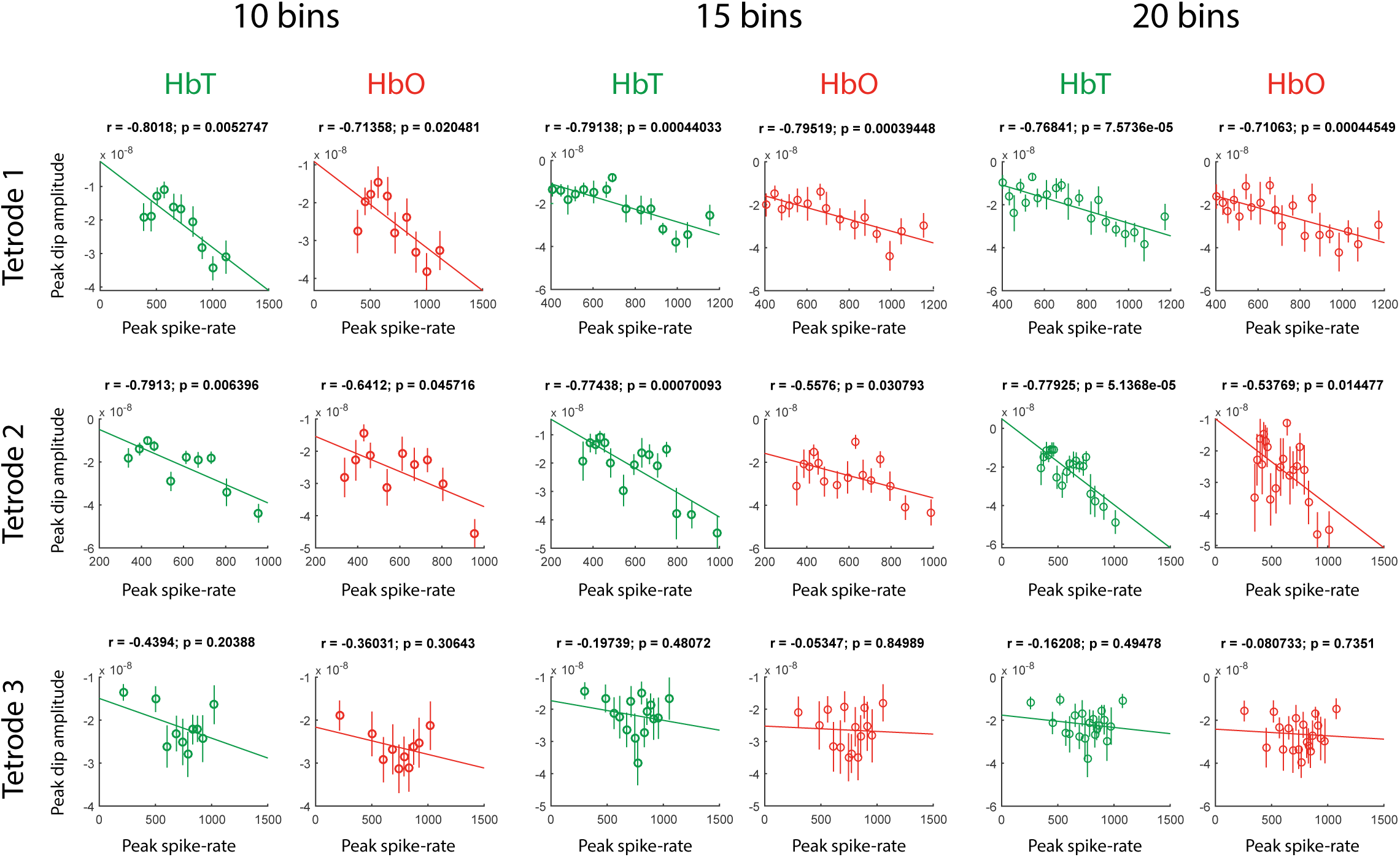
Correlations between peak amplitude of inital dip and peak spike-rates across tetrodes. Instead of using the mean slope between 0-1s, we used the peak dip amplitude (minimum signal value between 0 and 3s) for HbT and HbO and correlated it with the peak spike-rate. We found that the correlations of peak dip amplitude with peak spike-rates were strongest on the tetrode closest to the emitter, and decreased with increasing distance of tetrode from emitter (see figure 2H for comparison). These results were independent of the number of bins the trials were divided into. These results demonstrate that both HbO and HbT dip amplitudes can be used a proxy for underlying spiking activity.

**Figure S3.**
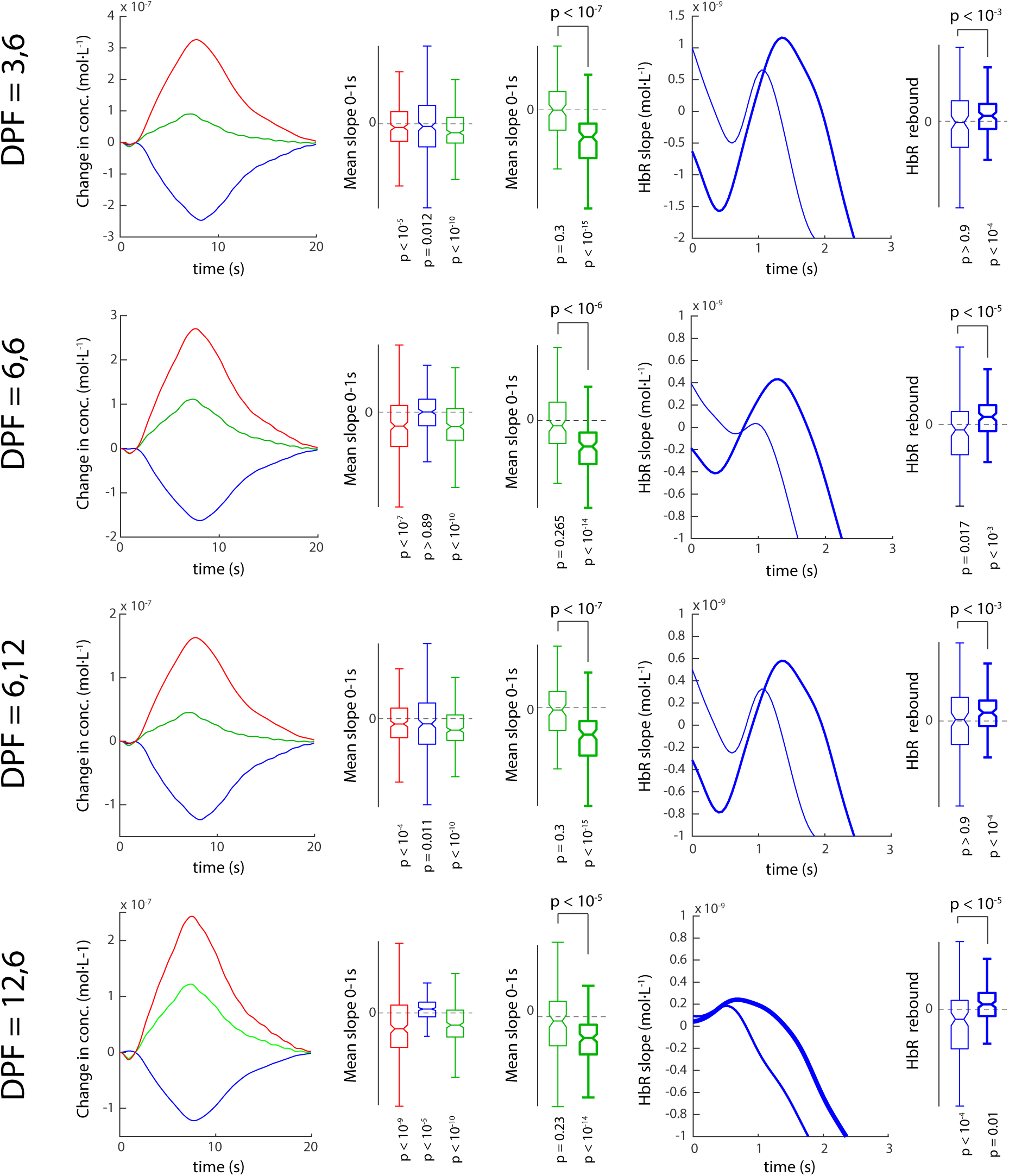
Verification of results across various differential path length factors (DPF). The conversion of optical density to concentration may vary based on the choice of DPF^1^. To ensure that our results did not depend on the DPF, we used multiple combinations of physiologically relevant DPF as reported earlier^1^ (the pair of values represent the DPFs at 760 and 850nm resp.) and found that although different DPFs lead to a change in the amplitude of the signals, the overall results were not affected. For each combination of DPFs, we observed strong dips in the HbO and HbT signals. We also observed an increase in the dip strength for HbT during high spiking trials. Low spiking trials failed to elicit a significant dip. The HbR signal also elicited a strong dip and rebound modulation in each case. For each combination of DPFs, the results were nearly identical and individually significant. Interestingly, the DFP combination 12,6 revealed increases in HbR within the 0-2s of trial onset. However, there was no difference in HbR concentration in high-spiking vs low-spiking trials (p>0.4; n=130/group, Wilcoxon rank-sum test). Furthermore, the strongest difference between high and low-spiking was still oberved in the HbT-dip (p<10^−5^) than HbO-dip (p<10^−3^). This reaffirms previous results^1^ that, at least in the primary visual cortex, although the choice of DPF might alter the amplitude of the calculated change in concentrations of HbO, HbR and HbT, it does not alter their relationship to underlying spiking activity. Notches represent 95% confidence intervals.

**Figure S4.**
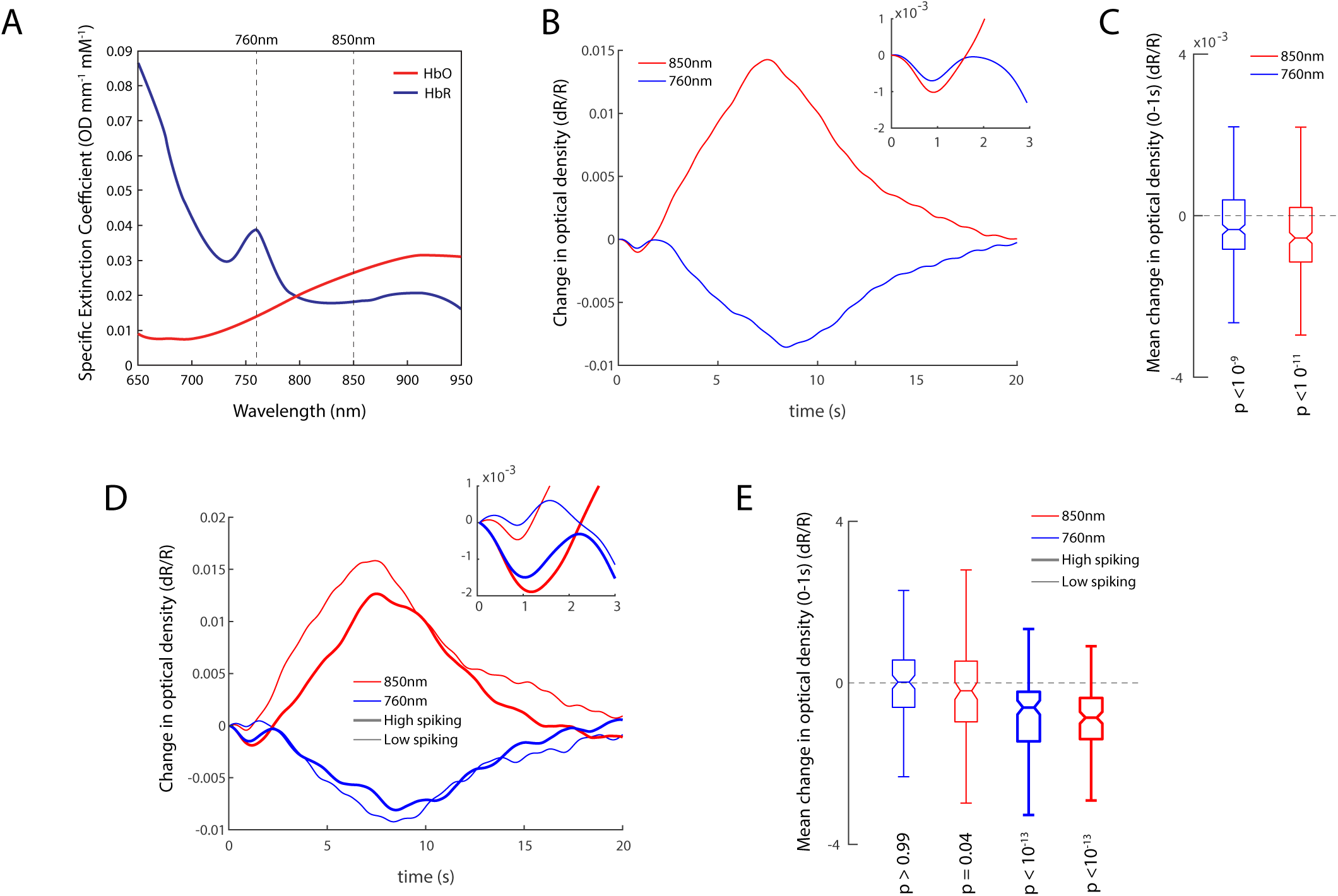
Changes in optical density for both imaging wavelenghts (760nm and 850nm) reveal an early decrease in chromophore concentration. A) Specific extinction coefficients for HbO and HbR as a function of wavelength. Although both HbO and HbR absorb light at both wavelengths, 760nm is more sensitive to HbR changes whereas 850nm is more sensitive to HbO changes. Changes in the observed optical density are proportional to changes in the chromatophores according to Beer’s law. B) Mean changes in the optical densities at 760nm and 850nm across all 260 trials. Inset represents the changes between 0-3s. A decrease in optical density for both wavelenghts can be observed. C) Mean change in optical density between 0 and 1s for 760 and 850nm. The distributions are significanlty less than zero. D) Optical density changes for trials with high and low spiking activity. Inset represents the changes between 0-3s. Larger decreases in optical density are observed for the high spiking trials. E) Mean changes in optical density between 0-1s reveal significant decreases in optical density for high-spiking trials. The decrease in optical density for both chromatophores implies a decrease in both HbO and HbR concentrations. Since HbT is the sum of changes in HbO and HbR, simultaneous decreases in HbO and HbR would lead to larger decreases in HbT when changes in optical density are converted to changes in concentration. Surprisingly, we found mildly significant decreases in the optical density at 850nm for low-spiking trials. However, after conversion to concentration changes, none of the traces (HbO, HbR or HbT) revealed significant changes between 0-1s.

**Figure S5.**
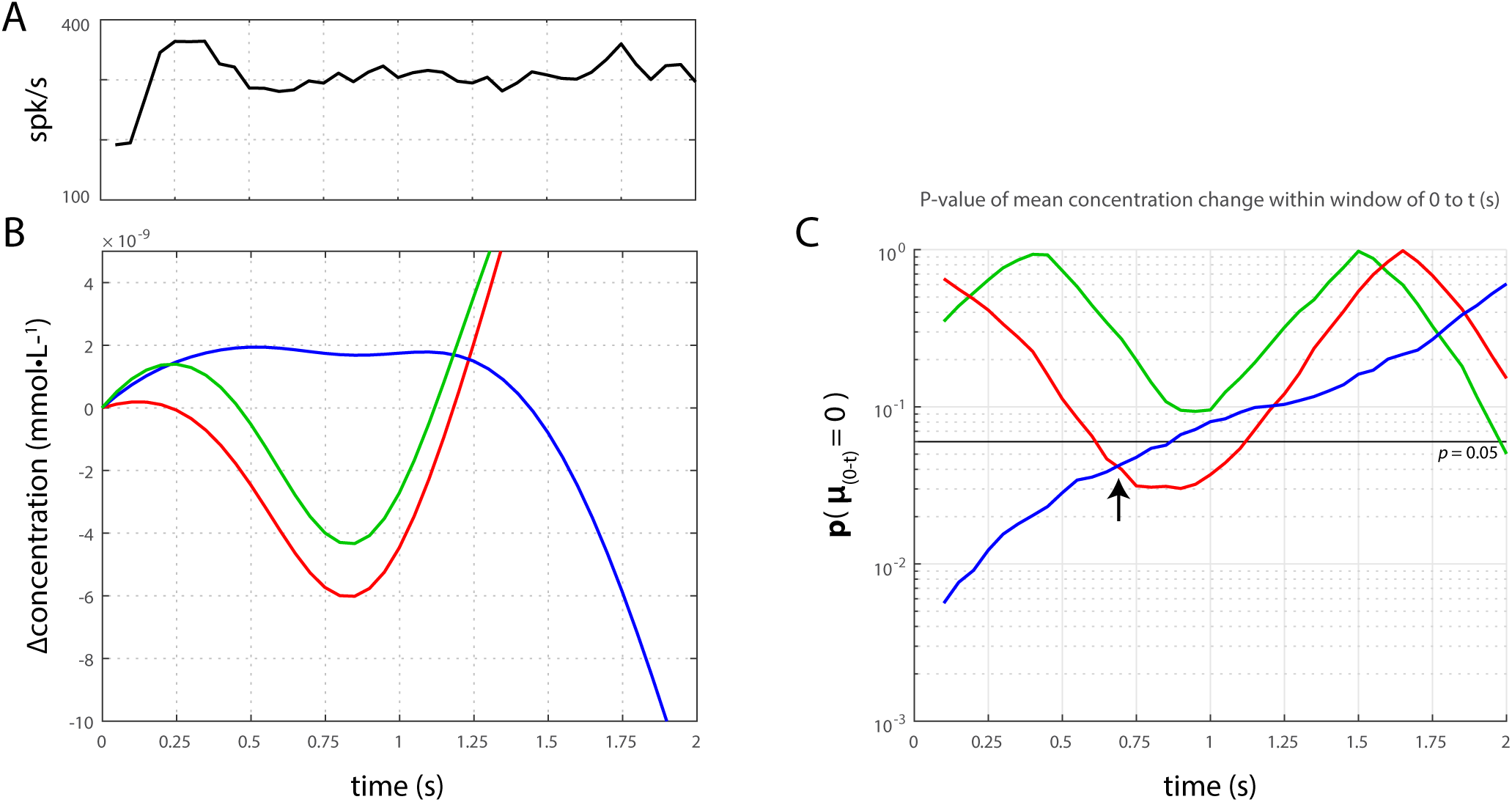
Further analysis of low-spiking trials reveals a significant increase in HbR and decrease in HbO concentrations within the first 750ms. We further analyzed trials where the peak spike-rate was less than the median value of all trials (A,see also Figure 2A). Although these trials belonged to the lower half of the distribution, they still had significantly high peak spike-rates and stimulus induced spike-rate moduations (see Fig. 2D). We obtained the mean HbO, HbR and HbT traces from these trials (B).Increases in HbR traces and decreases in the HbO traces are clearly observable within these trials. We determined the temporal window within which these changes were significant, by incrementing the width of the window from 0 to 2s (**C**, p<0.05 based on the Wilcoxon signed-rank test). We found the changes to be significant for HbO and HbR within 0-0.7s of stimulus onset (arrow). We failed to find significant changes for HbT for the same period (**C**, green trace). These findings demonstrate that during conditions of low spiking,the initial-dip consists of increases in HbR and decreases in HbO, and the HbT dip is absent. This suggets that the HbT dip is only induced
during conditions of strong bursts in spiking activity.

**Supplementary table 1.** Hemodynamic initial dips during spontaneous and stimulus-induced activity.

| dataset | parameter | condition | mean | SEM | p-value* | Sample-size |
| --- | --- | --- | --- | --- | --- | --- |
| Stimulus-induced activity | Peak spike-rate (spk/s) | all trials | 705.85 | 14.7 | $<10^{-10}$ | 260 |
| | | high-spiking | 899.96 | 12.89 | $<10^{-10}$ | 122 |
| | | Low-spiking | 497.28 | 8.35 | $<10^{-10}$ | 125 |
| | HbO dip ($\mu\text{mol}\cdot\text{L}^{-1}$) | All trials | -0.52 | 0.12 | $<10^{-7}$ | 260 |
| | | high-spiking | -0.87 | 0.146 | $<10^{-11}$ | 122 |
|  |  | Low-spiking | -0.22 | 0.188 | 0.21 | 125 |
| | HbR dip ($\mu\text{mol}\cdot\text{L}^{-1}$) | All trials | -0.04 | 0.078 | 0.89 | 260 |
|  |  | high-spiking | -0.2 | 0.089 | 0.0076 | 122 |
|  |  | Low-spiking | 0.09 | 0.12 | 0.18 | 125 |
| | HbT dip ($\mu\text{mol}\cdot\text{L}^{-1}$) | All trials | -0.56 | 0.11 | $<10^{-10}$ | 260 |
| | | High-spiking | -1.08 | 0.112 | $<10^{-14}$ | 122 |
|  |  | Low-spiking | -0.13 | 0.179 | 0.287 | 125 |
| | HbR buildup ($\mu\text{mol}\cdot\text{L}^{-1}$) | All trials | -0.041 | 0.805 | 0.634 | 260 |
| | | High-spiking | 0.236 | 0.093 | $<10^{-3}$ | 122 |
|  |  | Low-spiking | -0.365 | 0.133 | 0.007 | 125 |
|  | HbR- modulation (epoch II – epoch I) | All trials | 0.029 | 0.095 | 0.135 | 260 |
| | | High-spiking | 0.534 | 0.077 | $<10^{-10}$ | 122 |
| | | Low-spiking | -0.481 | 0.165 | $<10^{-3}$ | 125 |
|  | HbO/HbT dip (%) | High-spiking | 50.36 | 17 | - | 122 |
| Spontaneous activity | Total spike count | All runs | $1.56 \times 10^5$ | $1.03 \times 10^4$ | $<10^{-3}$ | 16 |
| | | High-spiking | $1.86 \times 10^5$ | $5.81 \times 10^3$ | 0.0078 | 8 |
| | | Low-spiking | $1.27 \times 10^4$ | $1.36 \times 10^4$ | 0.0078 | 8 |
| | HbO dip (a.u.) | All runs | -0.034 | 0.005 | $<10^{-6}$ | 48 |
| | | High-spiking | -0.051 | 0.012 | $<10^{-3}$ | 80 |
|  |  | Low-spiking | -0.036 | 0.015 | 0.06 | 80 |
|  | HbR dip (a.u.) | All runs | -0.009 | 0.005 | 0.1 | 48 |
|  |  | High-spiking | -0.032 | 0.014 | 0.0041 | 80 |
|  |  | Low-spiking | -0.001 | 0.016 | 0.9 | 80 |
| | HbT dip (a.u.) | All runs | -0.043 | 0.007 | $<10^{-6}$ | 48 |
| | | High-spiking | -0.07 | 0.016 | $<10^{-5}$ | 80 |
|  |  | Low-spiking | -0.037 | 0.018 | 0.122 | 80 |
|  | HbR buildup (a.u.) | High-spiking | 0.04 | 0.0136 | 0.011 | 80 |
|  |  | Low-spiking | 0.039 | 0.012 | 0.049 | 80 |
|  | HbR- modulation | High-spiking | 0.072 | 0.024 | 0.0017† | 80 |
|  |  | Low-spiking | 0.041 | 0.031 | 0.16† | 80 |
|  | HbO/HbT dip (%) | High-spiking | 58.9 | 32 | - | 48 |

\* Wilcoxon's two-tailed sign-rank test
† Wilcoxon's one-tailed sign-rank test

## Footnotes

1. Jasdzewski, G., Strangman, G., Wagner, J., Kwong, K.K., Poldrack, R.A. and Boas, D.A., 2003. Differences in the hemodynamic response to event-related motor and visual paradigms as measured by near-infrared spectroscopy. Neuroimage, 20(1), pp.479-488.

